# Establishing a causal role for medial prefrontal cortex in reality monitoring

**DOI:** 10.1101/436907

**Authors:** Karuna Subramaniam, Hardik Kothare, Leighton B. Hinkley, Phiroz Tarapore, Srikantan S. Nagarajan

## Abstract

Reality monitoring is defined as the ability to distinguish internally self-generated information from externally-derived information. Functional imaging studies have consistently found that the medial prefrontal cortex (mPFC) is a key brain region subserving reality monitoring. The aim of this study was to determine a causal role for mPFC in reality monitoring using navigated repetitive transcranial magnetic stimulation (nrTMS). In a subject-blinded sham-controlled crossover design, healthy individuals received either active or sham nrTMS targeting mPFC. Active modulation of mPFC using nrTMS at a frequency of 10 Hz, significantly improved identification of both self-generated and externally-derived information during reality monitoring, when compared to sham or baseline. Targeted excitatory modulation of mPFC also improved positive mood ratings, reduced negative mood ratings and increased overall alertness/arousal. These results establish optimal nrTMS dosing parameters that maximized tolerability/comfort and induced significant neuromodulatory effects in the mPFC target. Importantly, this is a proof-of-concept study that establishes the mPFC as a novel brain target that can be stimulated with nrTMS to causally impact both mood and higher-order reality monitoring.

---

Reality monitoring is defined as the ability to distinguish internally self-generated information from externally-derived information^1-4^. Reality monitoring is particularly relevant for patients with schizophrenia who suffer from cardinal impairments of the self, which directly contribute to their psychotic symptoms of delusions and hallucinations (indicating their break with reality)^5, 6^. We and others have recently shown using functional magnetic resonance imaging (fMRI) that the medial prefrontal cortex (mPFC) is a key brain region subserving reality monitoring, which has also been shown to be activated specifically during memory retrieval of self-generated information (i.e., recalling one’s own internal thoughts and actions)^3, 4, 7^.

Strong convergent evidence across lesion studies, meta-analysis of voxelbased morphometry (VBM) studies, deep brain stimulation and neuroimaging studies, all strongly suggest that mPFC is also a critical node for mediating mood state. For example, Downar and Daskalakis^8^ reviewed evidence across distinct and divergent methodologies to show that mPFC mediates mood enhancement in that: (i) lesions to the mPFC induced 80% risk of severe depressive symptomatology^9^; (ii) meta-analysis of VBM studies in Major Depressive Disorder (MDD) revealed the most reductions in volume were found in mPFC^10^; and (iii) DBS-induced inhibition of mPFC produced intense, dysphoria in a patient with remitted major depression ^11^.

We and others have also demonstrated that the mPFC plays a critical function in mediating interactions between mood and higher-order cognition during reality-monitoring^7, 12^. These prior correlative imaging studies which show that mPFC supports both reality monitoring and mood enhancement underscore the critical need to investigate whether mPFC can *causally* impact reality monitoring decision-making and mood. In this study, for the first time we use neurostimulation in the form of navigated repetitive transcranial magnetic stimulation (nrTMS) to establish the causal role of a novel neural target in the mPFC on impacting higher-order reality monitoring and mood in healthy controls (HC).

Repetitive transcranial magnetic stimulation (rTMS) has shown to be a robust neural plasticity-inducing technique that enables alterations of neuronal activity both in stimulated and remote areas via trans-synaptic neural plasticity^13, 14^. Lower frequencies of TMS at ~1 Hz typically produce inhibitory modulation effects, whereas TMS at more than 5 Hz are thought to generally produce excitatory modulation of the underlying cortical region being targeted^15^.

Although the first human studies of rTMS took place nearly 30 years ago^13, 16^, of all the stimulation targets tested to treat mood disorders and improve mood, the *medial* prefrontal cortex has received the least attention^8^. This is because the original applications of high-frequency TMS and the most widely used (and only) FDA-approved TMS therapeutic protocol have been applied to the dorsolateral prefrontal cortex, rather than the mPFC^17^. In these therapeutic TMS protocols, patients with Major Depressive Disorder (MDD) are predominantly treated with high-frequency stimulation in the dorsal lateral prefrontal cortex (DLPFC) of between 10-20 Hz for over 30 minutes^17, 18^. Given that the optimal parameters of stimulation are largely unknown particularly with regard to the mPFC, in the current study we test shorter protocols in HC that will enable greater affordability and broader transdiagnostic implementation across a range of disorders, such as in patients with mood and psychosis-spectrum disorders.

Here we apply high-frequency (10 Hz) nrTMS targeting mPFC to increase its activity^19^ (**Fig. 1A**). Targeting of mPFC was based on coordinates derived from our prior imaging studies on reality monitoring^3, 20^ (**Fig. 1B**). Our primary hypothesis was that increased excitatory activity induced by high-frequency 10Hz nrTMS targeting mPFC when compared to baseline and sham, would result in improvement of higher-order reality monitoring (**Fig. 1C and 1D**). Evidence in support of this hypothesis would thus establish for the first time an intimate causal relationship between mPFC activity and reality monitoring. A second hypothesis we tested here was that high-frequency nrTMS targeting mPFC would improve mood state in HC.

**Fig. 1.**
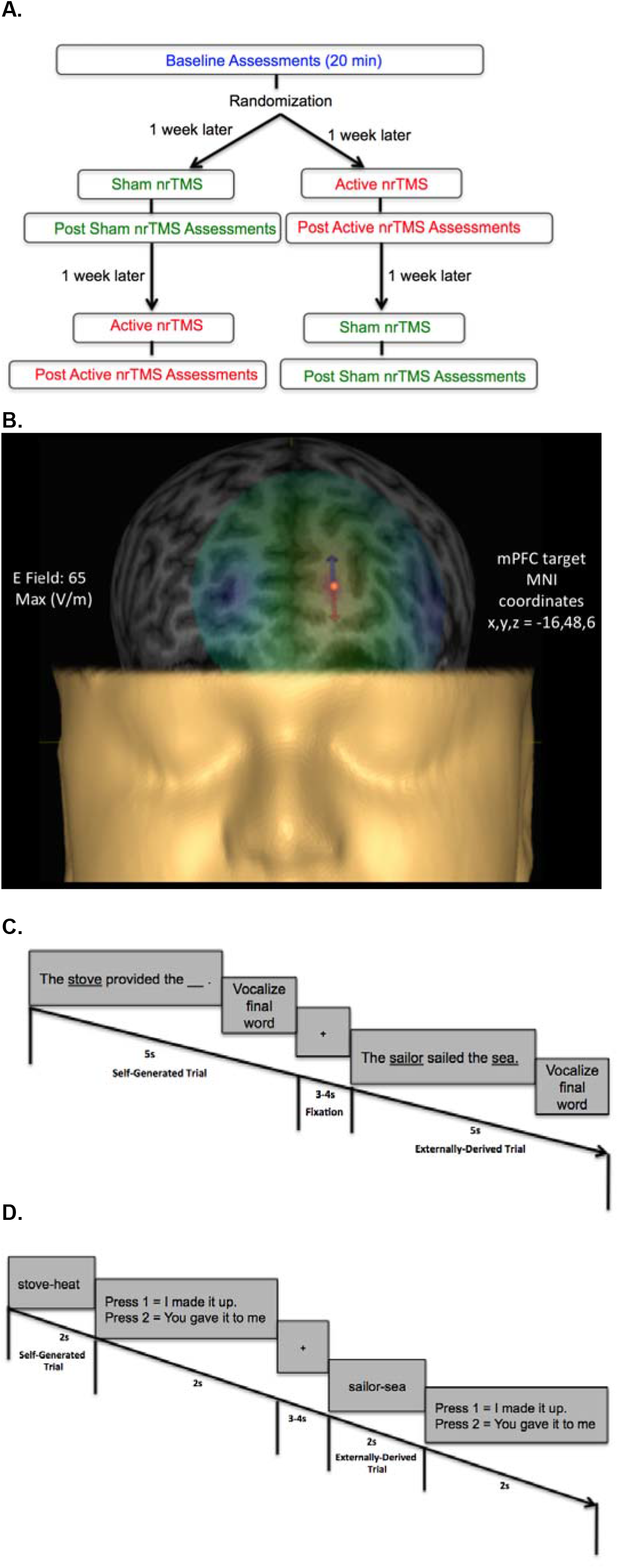
**A** Flow chart of randomized subject-blinded sham-controlled crossover nrTMS study trial. Subjects first completed baseline assessments, which included the reality monitoring encoding and retrieval tasks and mood and arousal ratings, for a total duration of 20 minutes. Subjects were then randomized to either the active nrTMS or sham nrTMS condition first; they completed the first nrTMS session a week after baseline assessments. After nrTMS application (duration=20 min), subjects immediately performed the post nrTMS reality monitoring assessments and mood/arousal ratings (duration=20 min). A week later, subjects completed the crossover (i.e., the second) nrTMS session and completed post nrTMS assessments (duration=20min) immediately after the nrTMS application. **B** A 3-D rendering of one subject’s head model is illustrated as an example, depicting the E-field strength in real-time when applying active high frequency 10Hz nrTMS to the mPFC target coordinates (x, y, z=-16,48,6), defined by peak reality monitoring activity in our prior neuroimaging studies^3, 20^. **C** Reality monitoring encoding task: Participants were presented with noun-verb-noun sentences in which the final word was either left blank for participants to generate themselves (e.g., *The <u>stove</u> provided the —*) or was externally-derived as it was provided by the experimenter (e.g., *The <u>sailor</u> sailed the <u>sea</u>*) For each sentence, participants were told to pay attention to the underlined words and to vocalize the final word of each sentence. **D** Reality monitoring retrieval task: Participants were randomly presented with the noun pairs from the sentences, and subjects had to identify with a button-press whether the second word was previously self-generated (e.g. stove-heat) or externally-derived (e.g. sailor-sea).

A final objective of the present study was to determine sufficient nrTMS dosage in HC, to test whether shorter protocols targeting the mPFC as a novel stimulation site in HC may be effective for improving both mood and higher-order cognition during reality monitoring. Prior studies have shown that high frequency 10Hz stimulation applied in 2s trains to the lateral PFC is beneficial to cognition^19^. Therefore, consistent with these studies^19^, here, we use similar parameters applied to the mPFC for the first time for a total duration of 20 minutes, which we hypothesized would yield beneficial mood and cognitive effects during a higher-order reality monitoring task assessed immediately after the nrTMS, but that would also meet safety and tolerability criteria in HC.

## Results

### MPFC modulation improves retrieval accuracy in reality monitoring

Active nrTMS targeting of mPFC at 10 Hz significantly improved overall reality monitoring performance, indexed by increased d-prime scores for overall accuracy, when compared to baseline (F[1,9]=12.32, p=.007) and sham conditions (F[1,8]=7.57, p=.03) (**Fig. 2A**). Improvements in retrieval accuracy, when compared to sham and baseline, were observed for both self-generated and externally-derived information (**Fig. 2B**). When compared to baseline, subjects accurately identified significantly more self-generated (F[1,9]=5.64, p=.04) and externally-derived information (F[1,9]=19.75, p=.002). When compared to the sham condition, subjects also accurately identified significantly more self-generated (F[1,8]=6.13, p=.04) and externally-derived information (F[1,8]=6.60, p=.03). There were no significant differences in reality monitoring performance between the sham nrTMS and baseline conditions or an order effect in nrTMS condition assignment on reality monitoring performance (all p’s > .20).

**Fig. 2.**
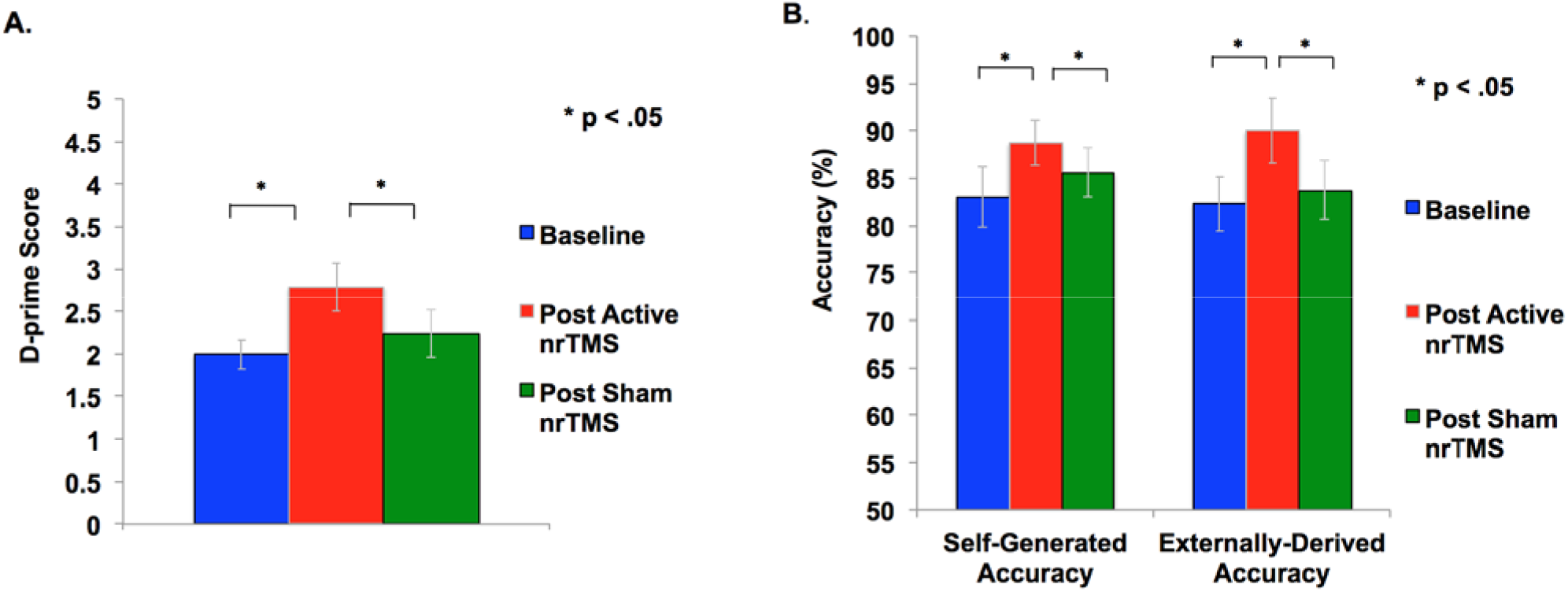
**A** Overall accuracy was computed as a d-prime score in each condition to differentiate sensitivity from response bias. Repeated-measures ANOVA revealed subjects had significant improvement in d-prime statistical scores for overall accurate identification of word items after active rTMS when compared to baseline or sham conditions. **B** Self-generated and externally-derived accuracy is illustrated as a percentage of the total number of self-generated and externally-presented trials in each condition (i.e., baseline, sham nrTMS, and active nrTMS), averaged across all subjects. Repeated-measures ANOVA revealed subjects had significant improvement in accurate retrieval of self-generated and externally-presented item-identification after active nrTMS when compared to baseline or sham nrTMS conditions.

### MPFC modulation improves mood

After active nrTMS of mPFC, subjects had significant improvements in positive mood (F[1,9]=5.26 p=.048) and significant reductions in their negative mood (F[1,9]=8.11 p=.02) when compared to baseline (**Fig. 3**). After active nrTMS when compared to sham, subjects showed marginal improvement in positive and negative mood states (F[1,8]=4.25, p=.07; F[1,8]=4.14, p=.07, respectively). They also showed significant improvements in overall alertness/arousal levels after active nrTMS when compared to baseline (F[1,9]= 7.36, p=.02) and sham conditions (F[1,8]=5.56, p=.046). There were no significant differences in mood/arousal levels between sham and baseline conditions, or an order effect in nrTMS condition assignment on mood/arousal levels (all p’s > .40).

**Fig. 3.**
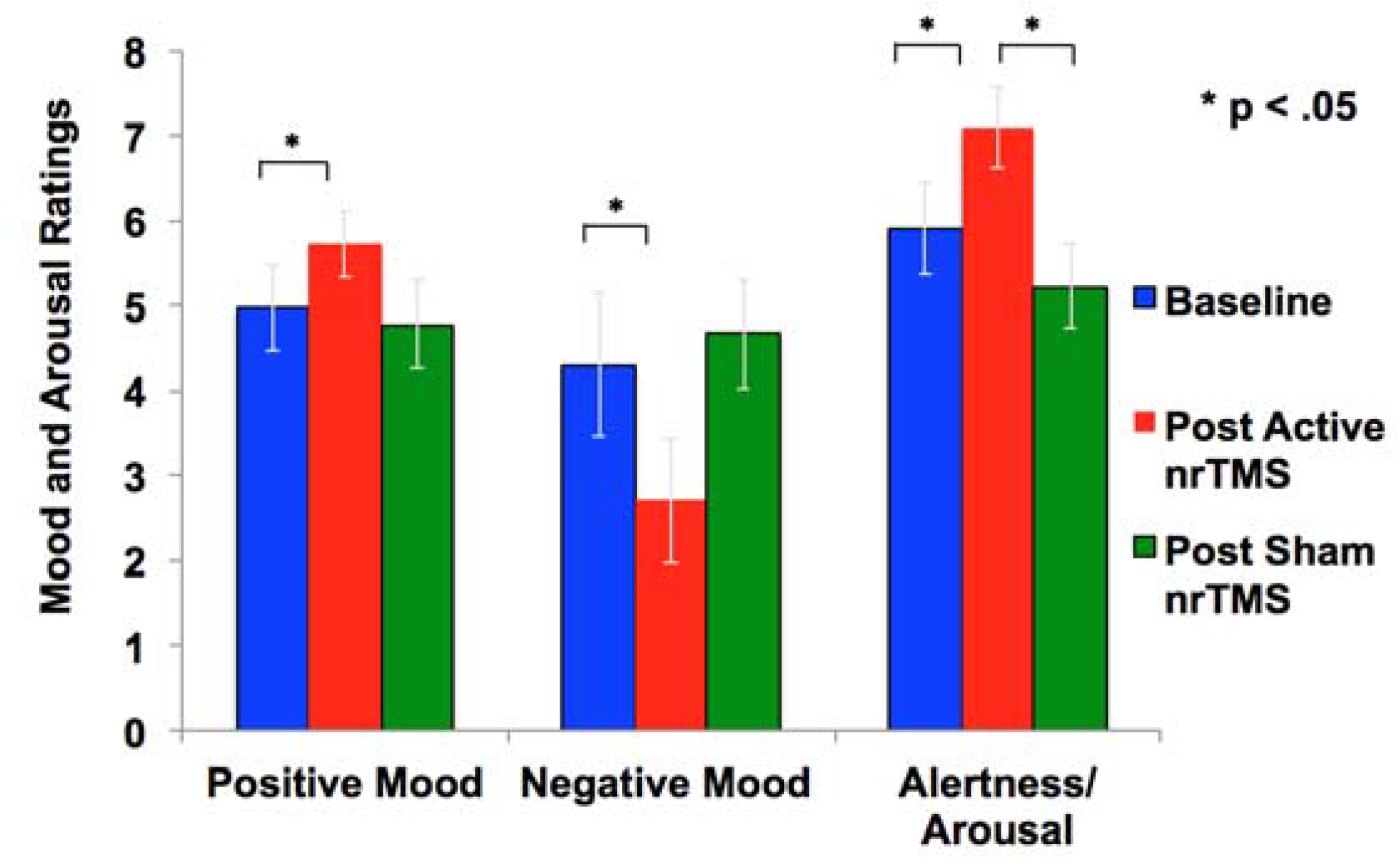
Mood and arousal/alertness in each condition (i.e., baseline, sham nrTMS, and active nrTMS) were rated on a visual analog scale from 0-9 (i.e., 0=Not at all; 9=Very high). Repeated-measures ANOVA revealed subjects had significantly increased positive mood, reduced negative mood and increased alertness/arousal levels after active nrTMS when compared to baseline assessments.

### Safety and tolerability of mPFC modulation with nrTMS

All HC (N=10) who completed active nrTMS reported that the active nrTMS was fairly highly tolerable and beneficial (6.15 and 6.20, respectively) on a scale of 0-9 (i.e., 0=Not at all; 9=Very high). No seizures or other serious adverse events occurred in any participant. Out of 11 subjects who were recruited, only one subject experienced referred pain at the back of her head (rather than the frontal region) after the first single nrTMS train for 2s duration. We ramped down the intensity after each single train in steps (i.e., to varying intensities at 100%, 90% and 80% of resting motor threshold (RMT)); however, this subject still experienced pain even at 80% of RMT. This subject reported no adverse events or side effects beyond the duration of each single train of nrTMS but was eliminated from the study and further analyses.

## Discussion

In the current subject-blinded sham-controlled crossover study, we found that the active high-frequency 10Hz nrTMS targeting this functionally localized focal peak of mPFC activity when compared to sham and baseline, significantly improved overall reality monitoring in HC, including accurate identification for both self-generated and externally-derived items. When compared to baseline assessments, active nrTMS significantly improved positive mood ratings, reduced negative mood ratings, and significantly improved alertness/arousal levels. Together, our results demonstrate for the first time that the mPFC represents a higher-order neural structure that causally mediates interactions between mood and higher-order reality monitoring that involves distinguishing between internal awareness of one’s own thoughts and actions from the outside external world. Consistent with our objectives, here we also established the sufficient nrTMS parameters that would yield significant mood and cognitive improvements in HC on our higher-order reality monitoring task but that would also meet safety and tolerability criteria, so that these same parameters could then be applied to a larger sample of HC and also be extended to patients with psychiatric disorders in future studies.

We have previously demonstrated in our fMRI studies that mPFC activation correlates with improved reality monitoring not only in HC, but also in psychiatric patient populations (i.e., patients with mood and psychotic disorders such as schizophrenia and schizoaffective disorders)^3, 7, 12^. Psychosis typically emerges during excessive pruning of excitatory pathways, leading to hypoactive aberrant networks in the PFC^21, 22^. Thus, these prior findings are particularly exciting, as they indicate that aberrations in mPFC activity are not immutably fixed even in chronically-ill patients with schizophrenia but that activity can be increased by behavioral interventions such that they became correlated with improvements in reality monitoring and real-world functioning^3, 20^. In other words, a serious cognitive reality monitoring deficit in patients with psychosis and its underlying neural dysfunction indexed by mPFC hypoactivity, can be improved by behavioral interventions and can improve quality of life even in patients with chronic psychosis^3^. Informed by our promising prior neuroimaging results, the nrTMS target we use here was, therefore, based on functional localization of mPFC activity mediating reality-monitoring in HC (x, y, z co-ordinates=-16,48,6), which also showed peak increase after cognitive training interventions in psychosis (x, y, z=-16,48,6)^3, 20^ (**Fig. S1A and S1B**). Together, our results demonstrate that the same region of mPFC that shows activation in HC, can also be activated to restore reality monitoring in psychosis^3^, indicating that the same mPFC target can be modulated with nrTMS in HC **(Fig. S1C)** as well as in patients with mood and psychotic disorders who suffer from severe reality monitoring impairments. This study, therefore, provides a promising neurobiological basis for a future precision medicine nrTMS approach. In particular, here, we localize the spatial aspects of nrTMS with regard to each participant’s neuroanatomy in which we adjust the scalp location of the stimulating magnet to the specific mPFC coordinates in which we observed peak reality monitoring activity in order to maximally engage these relevant mPFC networks mediating reality monitoring. In doing so, for the first time, we establish the powerful causal linkage of how nrTMS targeting mPFC as a novel neuromodulation target improves mood and reality monitoring in HC, thus providing a basis for future novel therapeutic interventions in psychosis in which reality monitoring is impaired.

The present paper also extends our previous findings in which we induced positive, neutral and negative mood states in healthy individuals while they performed fMRI to delineate how different mood states modulated subsequent reality monitoring performance. In these prior studies, we found that a positive mood significantly enhanced reality monitoring task performance in HC subjects via mPFC-activity enhancement^7, 12^. These previous findings demonstrate that mPFC plays a critical functional role in mood-cognition interactions particularly when people are in positive mood states, which subsequently helped them to improve their reality monitoring performance. Our data are also consistent with a recent study in which 10 Hz nrTMS targeting mPFC was shown to be effective in patients with Major Depressive Disorder (MDD), significantly reducing depression symptoms in half of the patients^23^. Additionally, another recent study in schizophrenia has shown that stimulation of prefrontal cortex with transcranial direct current stimulation improved emotion identification and general social cognition^24^. The present results, therefore, validate these prior studies and provide the first neuromodulatory demonstration that the mPFC, in particular, causally mediates mood-cognition interactions in that high frequency stimulation targeting mPFC not only significantly reduced negative mood and improved positive mood, but also induced significantly better reality monitoring performance when compared to baseline assessments.

The causal mechanisms as to precisely how high-frequency nrTMS targeting mPFC enhances mood and reality monitoring performance remain to be understood. We do not know the direction of causality, i.e., whether nrTMS first improved mood and overall alertness, which subsequently enhanced reality monitoring, or whether nrTMS improved information encoding and memory retrieval to enable better self-awareness, which then enabled people to attend better to their mood and alertness.

We did find that mPFC stimulation improved accurate memory retrieval of both self-generated and externally-derived information, contributing to significant improvement in overall reality monitoring performance. We know that TMS propagates trans-synaptically, and that stimulation of mPFC has direct anatomical and functional connections to the DLPFC^25, 26^, that is consistently activated during external working memory^27-31^. The original and most widely used applications of high-frequency TMS have been applied to DLPFC, which is strongly interconnected to mPFC^17^. These findings suggest that that high-frequency mPFC stimulation likely increased DLPFC activation which facilitated encoding and memory recall of externally-derived information through trans-synaptic long-term potentiation (LTP) ^14, 19^. LTP is observed in the enhancement of inter-neuronal signal transmission that outlasts the stimulation. LTP forms the basis of cellular mechanisms subserving learning and memory, based on the fact that memories are encoded by changes in synaptic strength^14, 19, 32^. An abundant of high frequency magnetic stimulation studies done in both humans and animals have demonstrated that TMS is a prevalent plasticity-inducing technique, which in humans is thought to be dependent on short-term changes of synaptic efficacy as well as on the activity of glutamate receptors and calcium signaling critical for overall learning (i.e., successful encoding of information) and memory retrieval^14, 33, 34^.

It must also be noted that there are likely some subjects who have anatomic variations in the locations of their trigeminal nerve bundles that render them unable to tolerate stimulation of this mPFC target, as was the case with the one subject who experienced referred pain at the back of her head. The benefit of nrTMS is that stimulation can be paused immediately for such subjects, unlike pharmaceutical medications that require longer durations and half-lives to be taken up by the whole brain and body, and which consequently result in much worse and long-lasting side-effects. By contrast, the nrTMS dosage can be modified in real-time to optimize patient comfort and ensure tolerability. In the most sensitive patients for whom no physiologically meaningful dose is tolerable, the subject may withdraw from treatment without having to experience protracted discomfort.

Further dose-response studies will be needed to examine duration and generalizability effects of nrTMS neuromodulation targeting mPFC. In future studies, we will use magnetoencephalography imaging (MEGI) before and after nrTMS to investigate the neural plasticity of how cortical networks are reconfigured after nrTMS-induced mPFC modulation. Improvements in reality monitoring that are related to enhanced mPFC activity will confirm the necessary/causal role of mPFC in the effects of stimulation in these future studies.

In sum, this is a proof-of-concept study that provides the first direct evidence to establish the mPFC as a novel neural target that can be modulated with nrTMS to causally impact both mood and higher-order reality monitoring. These pioneering findings shed new light on applications of nrTMS targeting the mPFC with several far-reaching implications. First, excitatory modulation of mPFC as a novel target in HC can be extended to patients with mood and psychosis-spectrum disorders in which reality monitoring is impaired, particularly in light of our prior neuroimaging findings that the same mPFC neural target is recruited in both HC and patients with psychosis-spectrum disorders^3^ (**Fig. S1A and S1B**). Second, this study also establishes the single-session nrTMS protocol parameters that meet safety and tolerability criteria and that are sufficient to induce significant neuromodulatory effects in this target. Finally, with our immediate post-stimulation assessments we are able to maximize LTP effects after only a single session of nrTMS. This may prove highly beneficial for future MEGI studies that explore timing and duration of cortical network oscillation reconfigurations and plasticity induced by nrTMS. In conclusion, these results provide a neurobiological basis for mPFC to be used as a potential transdiagnostic target for future precision medicine nrTMS therapeutic interventions that aim to improve mood and reality monitoring across disorders.

## Methods

### Participants

In the present study, we recruited 11 healthy participants (5 males, 6 females, mean age = 31.84, mean education = 18.95). This study was approved by the Internal Review Board (IRB) at the University of California San Francisco (UCSF) and all research was performed in accordance with IRB regulations at UCSF. Inclusion criteria for healthy participants were: no psychiatric/neurological disorders, including no personality disorders, no current or history of substance dependence or substance abuse, meets MRI criteria, good general physical health, age between 18 and 60 years, right-handed, and English as first language. Subjects were informed that they were participating in a randomized subject-blinded sham-controlled nrTMS trial in which they would need to complete assessments (of ~20 min duration) at three time-points, and therefore needed to maintain as constant a state as possible when compared to the baseline assessments (in terms of amount of sleep, caffeine, exercise, mood, and alcohol intake). Only the person administering nrTMS was aware of whether each participant received active nrTMS or sham nrTMS and was not allowed to discuss randomization with the subject (for complete details, see Active and Sham nrTMS section below). All participants gave written informed consent and then completed the reality monitoring task and mood and alertness/arousal ratings at baseline (**Fig. 1A**). Participants were then randomly assigned to active nrTMS or to the sham nrTMS condition first in our subject-blinded crossover design, and then assigned to the other nrTMS condition (**Fig. 1A**). All post nrTMS assessments were conducted immediately after each nrTMS session. Order of the active and sham nrTMS was counter-balanced across subjects. One participant was unable to tolerate the active nrTMS due to referred pain at the back of her head, and another participant completed the active nrTMS but was unable to complete the sham nrTMS condition due to relocation to another city.

### Reality Monitoring Task

All subjects completed the reality monitoring task at baseline and immediately after each active nrTMS and sham nrTMS session. Reality monitoring requires that subjects make higher-order judgments about distinguishing whether information was previously self-generated or externally-derived. As described in previous experiments, the reality monitoring task consisted of an encoding phase and a memory retrieval phase^3^ (**Fig. 1C and 1D**). During encoding, participants were visually presented with semantically constrained sentences with the structure “noun-verb-noun,” presented in blocks of 20 trials per run. On alternating half of the sentences, the final word was either left blank for participants to generate themselves (e.g., *The <u>stove</u> provided the __*) or was externally-derived as it was provided by the experimenter (e.g., *The <u>sailor</u> sailed the <u>sea</u>*) For each sentence, participants were told to pay attention to the underlined words and to vocalize the final word of each sentence. Participants then completed the reality-monitoring retrieval task where they were randomly presented with the noun pairs from the sentences (e.g., *stove-heat*), and had to identify with a button-press whether the second word was previously self-generated or externally-derived (**Fig. 1C and 1D**). At each time point, the reality monitoring task consisted of different sets of matched semantically constrained sentences, based on our previous studies^3^. The number of correctly identified self-generated and externally-derived trials was computed for each participant during the retrieval phase.

### Behavioural Statistical Analyses

Signal detection theoretic d-prime analyses for the reality monitoring task were conducted on overall accuracy in identification of word items by calculating the hit rate and the false alarm rate for self-generated and externally-presented items, then converting each measure to z-scores, and subtracting the false alarm rate from the hit rate in order to differentiate sensitivity during accurate performance from response bias. Self-generated and externally-derived accuracy was computed as a percentage of the total number of self-generated and externally-presented trials in each condition (i.e., baseline, sham nrTMS, and active nrTMS) for each subject, and then averaged across all subjects. Repeated-measures ANOVAs were implemented to examine differences in overall d-prime accuracy, as well as for correctly identified self-generated and externally-derived information between active nrTMS, baseline and sham conditions. Outliers were defined as values above/below 2 standard deviations from the mean. We did not find any outliers in the behavioural data. Mean accuracy in each condition is illustrated for overall d-prime accuracy, and for correctly identified self-generated and externally-derived information, averaged across all participants (**Table 1**).

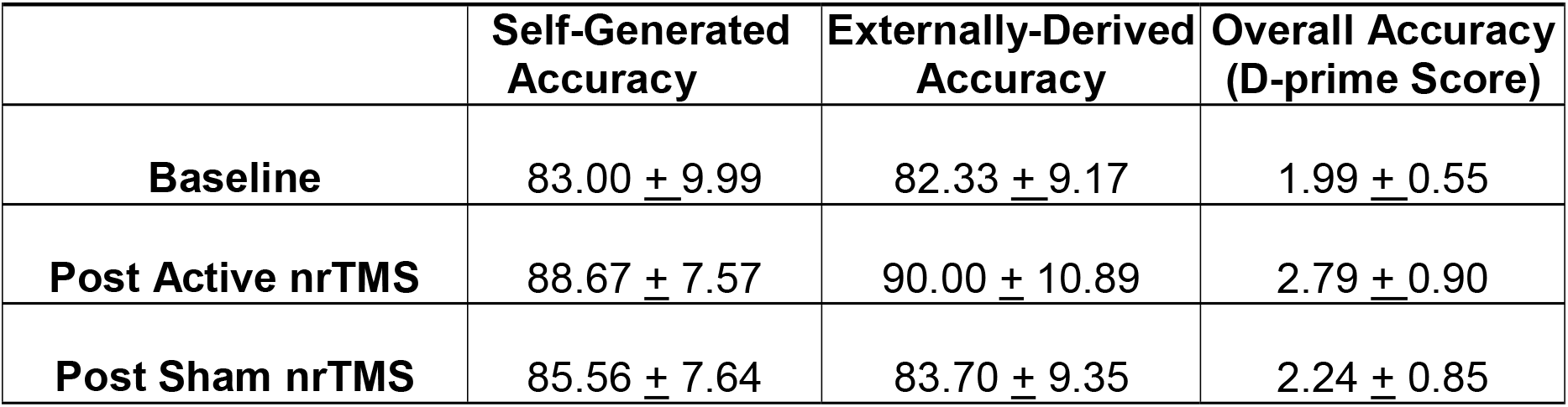
Table 1. Reality monitoring Accuracy

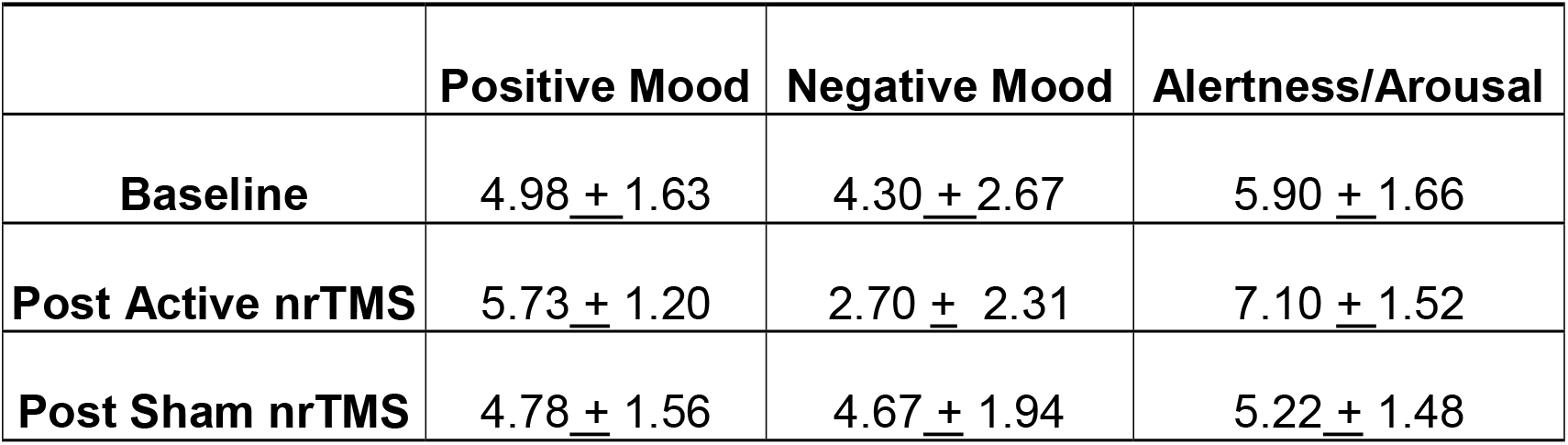
Table 2. Mood and Arousal Ratings

**Table 2. Mood and Arousal Ratings**
|  | **Positive Mood** | **Negative Mood** | **Alertness/Arousal** |
| --- | --- | --- | --- |
| **Baseline** | 4.98 ± 1.63 | 4.30 ± 2.67 | 5.90 ± 1.66 |
| **Post Active nrTMS** | 5.73 ± 1.20 | 2.70 ± 2.31 | 7.10 ± 1.52 |
| **Post Sham nrTMS** | 4.78 ± 1.56 | 4.67 ± 1.94 | 5.22 ± 1.48 |

We assessed positive mood, negative mood and arousal/alertness levels on a visual analog scale from 0 to 9 (i.e., 0=Not at all; 9=Very high) at baseline, and also immediately prior to and after each nrTMS active and sham session. Mood and alertness/arousal ratings for each condition are shown in **Table 2.**

### Safety and Tolerability of nrTMS

We first pilot tested 3 HC on 4 different nrTMS protocols prior to the start of the study to establish optimal dosage parameters that maximized tolerability/comfort in the present study. We tested each HC using the following nrTMS protocols, counterbalanced for each subject: (i) 20 Hz nrTMS for 2s trains at 110% of their resting motor threshold (RMT) with an ITI of 28s; (ii) 10 Hz nrTMS for 4s trains at 110% of their RMT with an ITI of 26s; (ii) 10 Hz nrTMS for 2s trains at 120% of their RMT with an ITI of 28s; (iv) 10 Hz nrTMS for 2s trains at 110% of their RMT with an ITI of 28s. The first protocol was least tolerable and most painful while the fourth protocol yielded highest tolerability ratings, and has also shown therapeutic efficacy in other studies^19^. In the present study, we therefore implemented the fourth nrTMS protocol for 20 minutes to examine its effects in healthy individuals during reality monitoring, given that nrTMS is known to induce long-term potentiation in neural activity in stimulated and remote brain areas expected to last beyond and for at least the length of the period of the nrTMS duration^14^ (i.e., here, we would expect nrTMS effects to last for at least 20 minutes after the TMS stimulation). Due to prior reports of painful ventromedial stimulation with 20Hz from our pilot test sample, we did not use higher frequencies than 10Hz or theta burst stimulation, or longer protocols and single train durations (>2s), which have a higher risk for pain and seizures. The online repetitive nrTMS parameters that we are using have also been deemed safe by the 2009 Consensus Guidelines^35^. Safety is defined by the absence of adverse events, such as seizures. As depicted in Rossi et al (2009)^35^, the maximum safe duration at 10Hz is >5 seconds. Thus, our exposure of 2 seconds at 10 Hz is well within the established range.

### Active and Sham nrTMS

After completing baseline assessments, participants were randomly assigned to active nrTMS or sham nrTMS condition first in a subject-blinded crossover design **(Fig. 1A)**. Targeting of nrTMS stimulation was carried out on the basis of a high-resolution anatomical T1-weighted MRI scan previously obtained for each subject. The nrTMS was delivered via state-of-the-art Nexstim Navigated Brain Simulation (NBS) (Nexstim Oy, Helsinki, Finland). This system integrates the TMS figure-8 coil with a software-based frameless stereotactic navigational system that allows for highly accurate cortical targeting that is individualized for each subject. Nexstim uses in-plane coil geometry to simulate the induced electric field and to make reliable predictions of electromagnetic field strength at the target location of neuronal activation^36^. The system calculates, in real-time, the strength of the electric field on the cortical target^37^. With the Nexstim system, we were thus able to calculate the electric field strength in real time as 65V/m for an individual subject (see **Fig. 1B**) when targeting the specific mPFC MNI coordinates (x=-16, y=48, z=6) that were based on our prior functional localization of mPFC peak activity mediating reality monitoring^3^. An integrated electromyography system allows for online detection of motor evoked potentials that result from TMS. This technology overcomes the problems of targeting and repeatability associated with traditional non-navigated TMS by allowing accurate spatial localization and field-strength calculations. Each participant completed 3 procedures: (I) MRI to Head Registration, which co-registers the patient’s anatomy to the navigational software; (II) Resting Hand Motor Threshold Determination, which maps out the hand motor area and determines each participant’s individual resting motor threshold; and (III) two nrTMS sessions, during which each participant received one active and one sham nrTMS which were separated by a week but conducted at roughly the same time of day for that subject.

#### (i) MRI to Head Registration

The nrTMS session began with aligning the MRI images to the subject’s head via the MRI to Head Registration process. Using the bridge of the nose and the crus of the helix of the ear, the 3D-locations of the landmarks (visible on both the subject’s MRI and the head) were measured using a digitizing pen with an optical tracking system. The optical tracking system used 2 cameras to triangulate the location in 3D space of infrared reflectors attached to the coil and subject’s head.

#### (ii) Resting Hand Motor Threshold Determination

Once the co-registration was completed, subjects completed single-pulse TMS mapping of the hand motor region. Resting motor threshold (RMT) was obtained by delivering single TMS pulses to the left motor cortex hand area. RMT was defined as the minimum stimulation intensity at which motor evoked potentials (thresholded at > 50 microvolts) were observed 50% of the time from surface muscle recordings from the first dorsal interosseous muscle of the right hand in half of the trials.

#### (iii) nrTMS Session

The target region for the nrTMS stimulation was the mPFC, located using MNI coordinates (x, y, z=-16,48,6) based on functional localization of mPFC peak activity mediating reality monitoring in healthy controls, which promisingly also showed peak activity increase (x, y, z=-16,48,6) in schizophrenia (SZ) patients after behavioral cognitive training interventions^3^. During the active nrTMS session, subjects received 800 pulses of 10 Hz nrTMS for 2s at 110% of their RMT with an ITI of 28s for a total duration of 20 minutes. nrTMS at this frequency and duration causes an increase in brain excitability over the stimulated area that lasted for the duration of the nrTMS session (at least 20 minutes)^19^. Pulse delivery was software controlled. The nrTMS application and immediate post nrTMS assessments altogether lasted approximately 40 minutes. Each participant also received sham stimulation in which the coil was placed over the same scalp region to target the same mPFC co-ordinates but with the stimulator output set to 1% of the RMT (producing an audible click). Only the person administering rTMS was aware of whether each participant received active rTMS or sham rTMS and was not allowed to discuss randomization with the subject. Order of the active and sham nrTMS was counter-balanced across subjects. We attained visual analog measures of tolerability and effectiveness of each rTMS session, as well as mood and arousal/alertness ratings before and after each nrTMS session rated on a scale from 0-9 (i.e., 0=Not at all; 9=Very high).

## Acknowledgments

This research was supported by the Brain and Behavior Research Foundation Young Investigator Award grant (NARSAD: 17680) and NIMH K01 grant (KO1MH82818) to Karuna Subramaniam, and the following NIH grants to Srikantan Nagarajan and John Houde (R01DC010145, R01DC013979, and R01NS100440).

## Author Contributions

KS recruited all the subjects and designed the reality monitoring experiment; acquired, analyzed and interpreted all the data; and wrote and edited the manuscript. HK helped with subject recruitment and acquisition of the nrTMS data. LH helped with subject recruitment and acquisition of the reality monitoring data. PT conducted the nrTMS sessions on all subjects, and edited the manuscript. SN edited the manuscript, and provided advice on the overall acquisition, analyses and interpretation of all the data.

